# Tobacco smoke alters the trajectory of lung adenocarcinoma evolution via effects on somatic selection and epistasis

**DOI:** 10.1101/2024.11.27.625765

**Authors:** Krishna Dasari, Jorge A. Alfaro-Murillo, Jeffrey P. Townsend

## Abstract

The mutagenic effects of tobacco smoke are well-reported. However, tobacco smoke also likely alters the somatic selective pressures that drive lung cancer evolution, and these altered pressures have never been quantified nor compared to the oncogenic effects of tobacco-induced mutagenesis. This comparison is necessary for prediction of targeted therapy efficacy given exposure history. Here, we estimated background mutation rates, somatic selection, and selective epistasis for 21 driver genes in ever- and never-smoker lung adenocarcinoma (ES-and NS-LUAD).

As expected, mutation rates were gene-specifically elevated in ES-LUAD. However, variant prevalence differences between ES- and NS-LUAD could not be explained by differential mutagenesis. *KRAS*, *KEAP1*, and *STK11* mutations were more strongly selected in ES-LUAD. *EGFR*, *PIK3CA*, and *SMAD4* mutations were more strongly selected in NS-LUAD, wherein *EGFR* mutations were associated with upregulation of epithelial-mesenchymal transition—a functional consequence specific to exposure history.

Epistasis was pervasive and context-dependent: ES-LUAD exhibited more synergy and less antagonism, implying a broader, more navigable adaptive landscape than NS-LUAD. We also performed the first systematic quantification of higher-order-than-pairwise epistasis, revealing sub-additive and emergent synergies that shape cancer evolution. Our results disambiguate the mutagenic and selective effects of tobacco smoke, demonstrating how environmental insults reshape cancer evolution and enabling prediction of therapeutic efficacy based on smoking history and somatic genotype.

**Author Summary:** Smoking is a well-known cause of lung cancer because it increases the number of mutations in lung cells. But tobacco smoke also causes inflammation and other physiological changes that might influence how tumors evolve. Using DNA from lung tumors, we asked whether these changes make certain mutations more helpful to cancer growth and thereby better targets for therapy. We also asked whether some mutations were more helpful to cancer growth in those who haven’t smoked. To do so, we estimated not only how often mutations occurred, but also how strongly they were favored during cancer development. Focusing on 21 key lung cancer genes, including *KRAS*, *EGFR,* and *TP53*, we found that mutations in some, such as *KRAS* and *KEAP1* were more strongly favored in smokers. Mutations in genes such as *EGFR* and *PIK3CA* were more strongly favored in non-smokers, where they were also linked to non-smoker-specific changes in gene activity. We also found that mutations in tumors interact differently depending on smoking history: smokers’ tumors showed more cooperation between mutations, which may help them evolve in more flexible ways. This study shows how smoking reshapes the genetic paths lung cancers take—and why a patient’s smoking history matters when choosing targeted treatments. Understanding these evolutionary patterns can help improve treatment decisions and lead to more personalized care for lung cancer patients.

## Introduction

Tobacco smoke is well known to cause cancer, in part through its mutagenic effects [1]. Carcinogen exposure from tobacco generates distinct patterns of mutations [2], a subset of which then contribute to oncogenesis [3–9]. In addition to this mutagenic effect, tobacco smoke induces physiological changes in the lung—inflammation, oxidative stress, immune dysregulation, and tissue damage [1,10–12]—that can alter the lung microenvironment and thereby the selective pressures acting on specific somatic mutations. These microenvironmental disruptions could modulate the fitness advantages conferred by specific mutations, thereby influencing their likelihood of contributing to cancer development. Indeed, several studies have suggested that, in principle, endogenous or exogenous alterations to the microenvironment caused by factors such as aging or tobacco exposure likely shape the trajectories of cancer evolution [13–17]. However, the extent to which such physiological effects alter the adaptive landscape of cancer has not previously been quantified. Therefore, quantification of the adaptive impacts of a highly disruptive exogenous insult such as tobacco smoke is crucial to understanding how environmental exposures shape the evolution of individual cancer evolutionary trajectories.

Among lung cancer subtypes, LUAD is the most common and accounts for the highest proportion of cases in never-smokers [18]. As of 2002, nearly 23% of LUADs in the United States occurred in never-smokers [19], a proportion that has been rising as cigarette usage declines [20]. Numerous studies have reported that LUAD exhibits distinct somatic variant profiles in never-smokers compared to ever-smokers, which has been taken as evidence of their divergent genetic vulnerabilities [21–25]. However, differences in variant prevalence alone are confounded as to whether they are a consequence of effects of tobacco-induced mutagenesis or a consequence of tobacco-altered selection dynamics. Therefore, quantification of the relative oncogenic roles of the mutagenic and physiological effects of smoking in LUAD will be increasingly crucial to determination of optimal use of targeted therapies for patients with and without a prior smoking history.

Computational models now enable inference of baseline somatic mutation rates from synonymous mutations observed in tumor genomes—implicitly [6] or explicitly [26] accounting for carcinogen-induced mutagenesis, such as that caused by tobacco smoke. Comparing these baseline mutation rates to the observed prevalence of nonsynonymous mutations enables estimation of somatic selection acting on specific substitutions [8,27,28]. This framework provides a foundation for disentangling the oncogenic contributions of the mutagenic and selective effects of smoking. Moreover, selection on each mutation is likely influenced not only by the microenvironmental effects of tobacco smoke, but also by pairwise [28–30] and higher-order [28] selective epistasis between mutations. In selective epistasis, mutations affect the selection operating on other mutations. These interactions between the selective effects of mutations shape the adaptive landscape of cancer evolution [5,28,31,32]. Understanding how tobacco smoke influences not just mutation and selection, but also epistasis, is essential for a complete picture of LUAD evolution.

Because knowledge of epistasis is crucial to inference of cancer-driving mutation modules and synthetic vulnerabilities, a number of algorithms have been developed that attempt to detect epistasis between mutations or sets of mutations by quantifying over- and under-representation [33,34] in the context of heterogeneous mutation rates [30,35–37]. However, with the exception of Coselens [30], these methods do not enable meaningful quantitative comparisons of the effects of mutations in distinct somatic genotypes. This quantification is a crucial step for prioritization and translation of knowledge of epistasis into effective therapeutic strategies. Moreover, all of these approaches are restricted to pairwise inferences [30,34] or to inference of discrete modules [33,35–37], and are not well suited to the quantitative inference of higher-order epistasis (involving more than two mutations). When present, higher-order epistasis substantially influences the adaptive landscape and shapes evolutionary trajectories [38–43], including the degree to which selective epistasis diminishes along the adaptive trajectory [44–47]. Systematic characterization of the presence and biological significance of higher-order epistasis to cancer evolution has been challenging. Recent theory facilitates understanding of higher-order epistasis in cancer by providing the selection strength of mutations for each somatic genotype [28]. This novel approach has not yet been applied in the context of environmental risk factors that could also affect selection, such as tobacco smoke. Accordingly, analysis of the effect of smoke on the adaptive landscape of LUAD will benefit from consideration of pairwise and higher-order epistatic effects and potential differences in epistasis between microenvironments.

To address these gaps, we quantified the mutagenic, selective, and epistatic effects of tobacco smoke on single-nucleotide variants (SNVs) in LUAD. We assembled a large dataset of lung adenocarcinoma tumor genomes and classified them by smoking status into ever-smoker (ES) and never-smoker (NS) groups. We estimated background mutation rates in each group and inferred the strength of somatic selection acting on driver mutations in each gene. We then investigated how selection pressures differed between ES- and NS-LUAD and associated these differences with differential gene expression. Finally, we extended our analysis to inference of selection in specific somatic genotypes, detecting and quantifying the extent of pairwise and higher-order epistasis. The degree of divergence in the mutational, selective, epistatic, and transcriptional landscapes of LUAD between ever-smokers and never-smokers enables novel insight into how environmental exposure shapes cancer evolution—identifying genotype- and environment-specific vulnerabilities that can inform personalized therapeutic strategies.

## Results

### Mutation rates and somatic selection pressures jointly shape the prevalence of driver variants in lung adenocarcinoma

To disentangle the relative contributions of mutation and selection to observed mutation frequencies in LUAD, we deconvolved the prevalence of somatic variants in our curated set of 21 well-supported driver genes into estimates of the background oncogenic mutation rates and scaled selection coefficients. Across all smoking-status-classified LUAD samples in our dataset (*n* = 1,722), *TP53* and *KRAS* were the most frequently mutated genes, followed distantly by *EGFR*, *KEAP1*, and *STK11* (**Fig. 1A**). Background oncogenic mutation rates for the majority of these genes spanned slightly more than an order of magnitude—between 7.6 × 10^−8^ and 8.7 × 10^−7^ (**Fig. 1B**). When compared to the observed variant prevalences, this variation in background oncogenic mutation rate frequently implies substantial and variable impacts of somatic selection on the prevalence of each variant.

**Figure 1.**
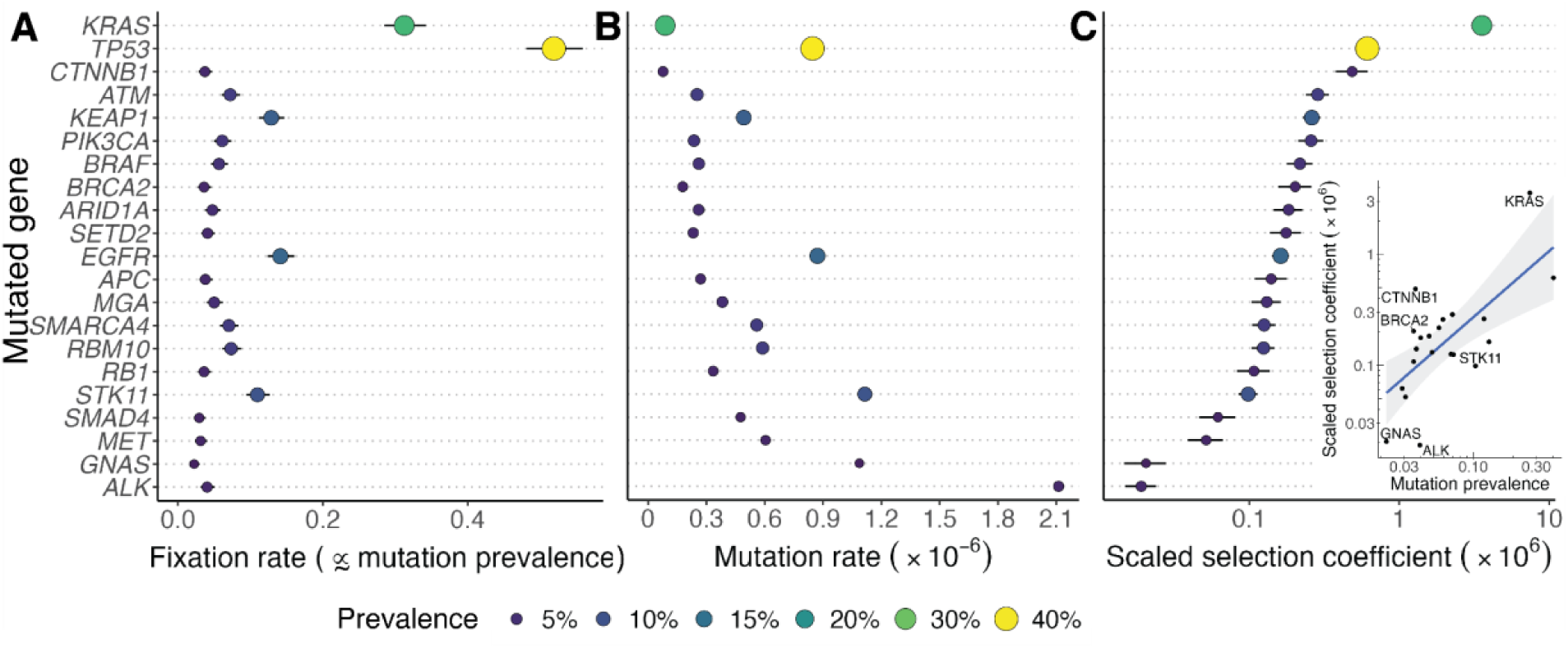
Fixation rates and scaled somatic selection coefficients for 21 known and potential driver mutations among 1,722 lung adenocarcinomas. **(A)** Poisson-corrected fixation rates (rates at which each mutation occurs and becomes fixed within the population). **(B)** Background rate at which oncogenic gene mutations occur. **(C)** Strength of somatic selection (log-scaled) experienced by oncogenic gene mutations. Inset: Correlation between the prevalence of an oncogenic mutation and the strength of selection acting on it (*P* < 0.001).

Scaled selection coefficients for oncogenic mutations in each gene exhibited a long-tailed distribution (**Fig. 1C**), with mutations in *KRAS* experiencing the strongest selection, followed distantly by mutations of *TP53* and *CTNNB1*, then mutations in *ATM*, *KEAP1*, and *PIK3CA*, and then other genes that experienced weaker but still substantial selection (**Fig. 1C**). Notably, *CTNNB1* mutations were present in only 3.7% of tumors. Nevertheless, knowledge of the low underlying oncogenic mutation rate of *CTNNB1* indicates that these infrequent mutations experienced stronger selection than did the more frequent mutations in *STK11, RBM10*, and *SMARCA4*. Generally, selection and mutation prevalence were correlated (*r* = 0.68, *P* < 0.001), but several discrepancies were present between the prevalence of a mutation and the relative intensity of selection it experienced (**Fig. 1C**, inset). These discrepancies highlight the importance of reporting selection as the effect size of mutations, rather than reporting solely prevalence, which is only moderately associated with strength of selection.

The oncogenic mutation rates and selection coefficients presented above were quantified based on all tumors, regardless of smoking status. However, both the mutagenic processes and selective advantages of mutations were plausibly influenced by the genotoxic and physiological effects of tobacco smoke. To assess these distinct effects, we stratified tumors by smoking status and performed separate analyses of somatic selection in ES- and NS-LUAD tumors.

### Mutation and selection differ substantially between ever-smoker and never-smoker LUAD

Stratification of LUAD tumors by smoking status based on mutational signature attributions and clinical annotation yielded 1,066 ES-LUADs and 656 NS-LUADs. In each group, we independently estimated oncogenic mutation rates and selection coefficients for the 21 well-established driver genes (**Fig. 2**). Both mutation rates and selection pressures differed markedly between ES- and NS-LUAD. Oncogenic mutation rates were elevated in ES-LUAD across all 21 genes. Increases ranged from modest (39%–43% for *RB1* and *CTNNB1*) to extreme (571–1474% for *EGFR*, *SMAD4*, *GNAS*, and *ALK*; **Fig. 2A inset**). These elevated mutation rates demonstrate the intense mutagenic effect of tobacco smoking on the peripheral tissue of the lung.

**Figure 2.**
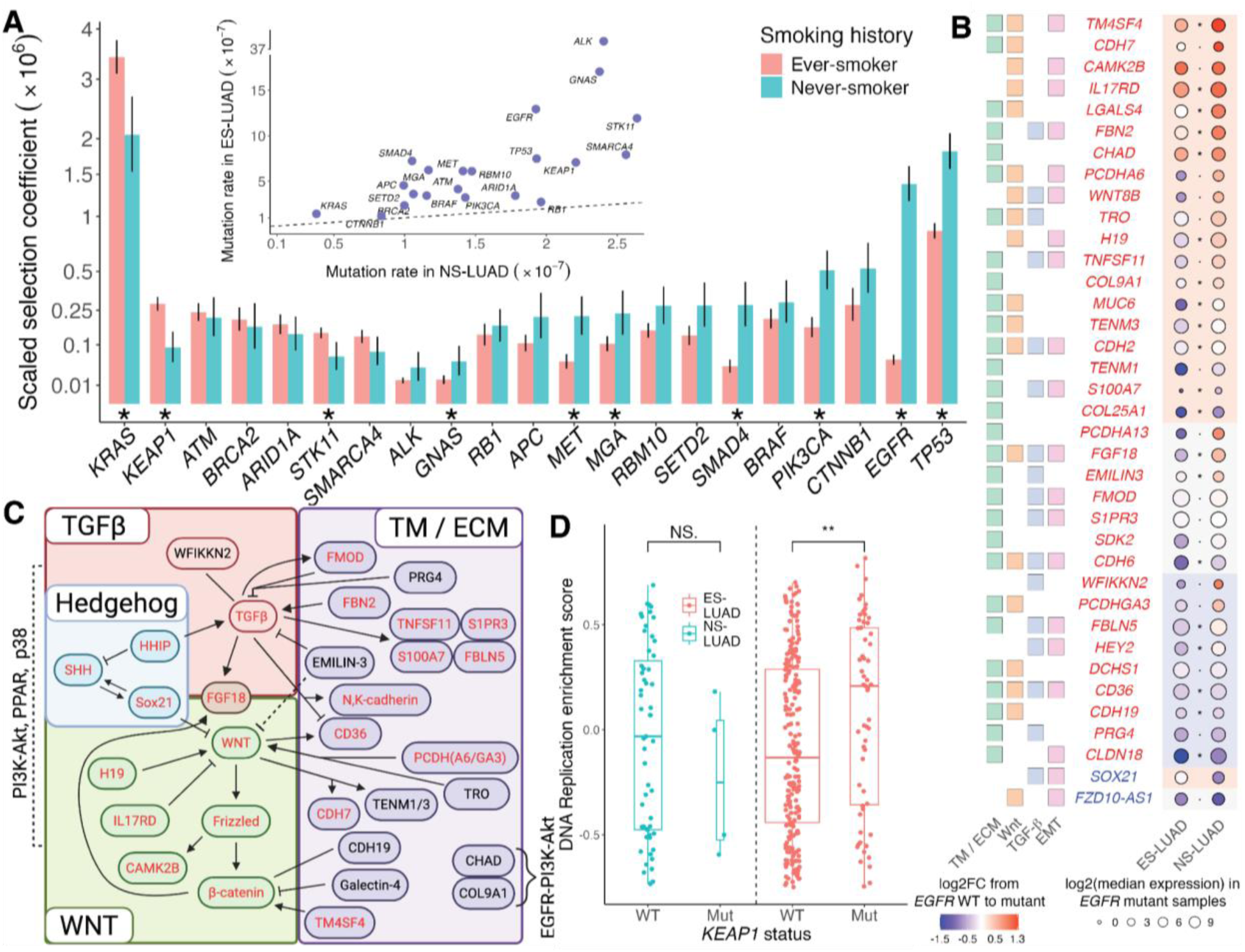
Mutational and adaptive landscapes of ever-smoker lung adenocarcinoma (ES-LUAD) and never-smoker lung adenocarcinoma (NS-LUAD). (**A**) Strengths of somatic selection (*y* axis is square-root scaled; *: non-overlapping 95% confidence intervals) for mutations of 21 known and potential driver genes in NS-LUAD (teal bars; *n* = 656) and ES-LUAD (salmon bars; *n* = 1,066), averaged across somatic genotypes. Inset: Background oncogenic mutation rates for these genes in ES-LUAD and NS-LUAD (dashed line: equality). (**B**) Differential expression of genes between *EGFR* wild-type (WT) and *EGFR* mutant samples in ES-LUAD (*n* = 273) and NS-LUAD (*n* = 59; log2FC: log_2_ of fold change in expression) for genes deemed to have a significant gene expression interaction between *EGFR* allelic status and smoking status. Color of each gene name denotes whether the change in expression was more positive (red) or more negative (blue) in NS-LUAD compared to ES-LUAD (*: *P* < 0.05, •: *P* < 0.1, *SOX21*: *P* = 0.116; background color: change in gene expression from normal (*n* = 33) to LUAD (*n* = 332) is increased [pink], decreased [light blue], or insignificant [gray]). Leftward squares designate whether the gene produces a transmembrane (TM) or extracellular matrix (ECM) protein as well as whether it facilitates or interacts with the Wnt, and TGF-β and the epithelial-mesenchymal transition (EMT) signaling pathways. (**C**) Interactions between proteins (red: implicated in EMT) whose *EGFR*-mutation-associated expression changes differ between ES-LUAD and NS-LUAD. (**D**) Sample-specific DNA replication expression scores for *KEAP1* WT and mutant ES- and NS-LUAD (**: *P* < 0.01).

We found that reported differences between ES- and NS-LUAD in the prevalences of key driver mutations [21,22,48] were often driven by differential selection, rather than solely by differential mutagenesis (**Fig. 2A**). Among the 21 driver genes analyzed, *TP53*, *EGFR*, *PIK3CA*, *SMAD4*, *MET*, *MGA*, and *GNAS* mutations experienced stronger positive selection in NS-LUAD. *STK11*, *KEAP1*, and *KRAS* mutations experienced stronger positive selection in ES-LUAD. *RB1*, *BRAF*, and *BRCA2* mutations experienced similar selection in both cancer types.

Most mutations that experienced significantly different selection between ES- and NS-LUAD were strongly selected in one context and weakly selected in the other, demonstrating pronounced context-dependence in LUAD evolution. In ES-LUAD, *KEAP1* mutations experienced the third strongest selection—after *TP53* and *KRAS*—and experienced over three times stronger selection than in NS-LUAD. Likewise, *STK11* mutations experienced over twice as much selection in ES-LUAD. Indeed, both *KEAP1* and *STK11* mutations are enriched in ever-smoker populations [23,49,50]. *EGFR* mutations exhibited the most dramatic difference: they were more than twenty times more strongly selected in NS- than ES-LUAD, and were also the third most strongly selected mutations in NS-LUAD after *TP53* and *KRAS*. *EGFR* mutations are much more common in NS-LUAD than in ES-LUAD [21,51]. Similarly, *PIK3CA*, *SMAD4*, and *MET* mutations experienced over three times stronger selection in NS-relative to ES-LUAD.

To investigate mechanisms underlying the differential selection of *EGFR* mutations, we assessed their differential transcriptional effects in ES- and NS-LUAD using TCGA expression data. In fewer than one percent of genes did the expression fold change from *EGFR* wild type samples compared to *EGFR* mutant samples differ significantly between ES- and NS-LUAD (**Fig. 2B**). However, several of these genes exhibited markedly stronger upregulation or downregulation in EGFR-mutant NS-LUAD compared to ES-LUAD. Genes that were upregulated were frequently components of Wnt, TGF-β, and Hedgehog signaling pathways or of the extracellular matrix (**Fig. 2B–C**). Indeed, in an analysis of gene sets upregulated in NS-versus ES-LUAD, the Hallmark gene set associated with epithelial-mesenchymal transition was the most enriched. On the other hand, *KEAP1* mutations were associated with a substantially higher DNA replication enrichment score in ES-LUAD but not in NS-LUAD (**Fig. 2D**). These examples relate microenvironmental context to modifications of the downstream effects of driver mutations and their accompanying selective advantage.

Extending our analysis from 21 to 1,200 LUAD-associated genes revealed a striking asymmetry in selection by smoking status: only six genes—*KRAS*, *KEAP1*, *STK11*, *PBRM1*, *DNMT3B,* and *CDH1—*were significantly more strongly selected in ES- than NS-LUAD, whereas 87 experienced stronger selection in NS-LUAD (**Fig. S1**). Among the mutations most strongly favored in NS-LUAD were those in *EGFR*, *SMAD4*, *SMAD2*, *PTEN*, *PIK3R1, PAK1*, *FGF3*, *GATA3*, *JUN*, *CTNNA2*, and *TLR4*. Mutations often experienced weaker selection in ES-LUAD, which is more heavily mutated and frequently possesses multiple known drivers, than in NS-LUAD, which often features a single known driver (**Fig. S1**).

Overall, somatic mutations often experienced substantially altered selection in the tobacco-smoke-altered ES-LUAD compared to NS-LUAD, contributing to differential evolution of the two cancers. Along with the somatic environment, the somatic genotype can also induce differential selection on specific mutations: effects of each mutation can be analyzed across somatic genotypes to reveal selective epistasis that shapes the evolutionary trajectories of tumors in ever-smokers and never-smokers. Therefore, we assessed the influence of pairwise selective epistasis.

### Pairwise epistasis shapes the adaptive landscape of ES-LUAD and NS-LUAD

To assess how frequently and how strongly the evolutionary trajectory of LUAD is affected by pairwise epistasis, we quantified the effects of driver mutations both in isolation (in tumors that were wild type at a paired gene) and in the presence of each paired gene mutation. For ES- and NS-LUAD, we evaluated all 420 possible gene pairs among the 21 driver genes: pairwise interactions were identified that modulate the selective advantage of mutations, and did so in a smoking-dependent manner.

#### Mutation of *TP53* followed by *RB1* is the most likely trajectory for ES-LUAD and NS-LUAD

To investigate directional epistasis between two common LUAD drivers, *TP53* and *RB1*, we first stratified tumors by smoking status and somatic genotype at these two loci, and then estimated the fixation rates of each mutation from each somatic genotype (**Fig. 3A**). In ES-LUAD, *TP53* mutations exhibited a high fixation rate when *RB1* was wild-type but were inferred to exhibit a negligible fixation rate when *RB1* was mutated (***λ*, Fig. 3B**). Conversely, *RB1* mutations exhibited a low fixation rate when *TP53* was wild-type but exhibited a high fixation rate when *TP53* was mutated. These asymmetries indicate the most likely evolutionary sequence in ES-LUAD: mutation of *TP53* followed by mutation of *RB1*. In wild-type tumors, the mutation rate of TP53 was over double the mutation rate of *RB1* (*µ*_WT→TP53_ *=* 7.4 × 10^−7^, *µ*_WT→RB1_ *=* 2.0 × 10^−7^); the same was true of mutant tumors (*µ*_RB1→RB1+TP53_ *=* 8.9 × 10^−7^, *µ*_TP53→RB1+TP53_ *=* 3.6 × 10^−7^; **Fig. 3B**). Accordingly, *RB1* variants in a *TP53*-mutant genotype must have experienced comparable selection to *TP53*-variants in a wild-type genotype (*γ*, **Fig. 3B**), despite the substantially lower fixation rate of the *RB1* variants. This pattern reflects synergistic epistasis: prior mutation of *TP53* increases selection for *RB1*. In contrast, the reverse order exhibited antagonistic epistasis—*TP53* was less strongly selected in the presence of *RB1* mutation—though this last effect did not achieve statistical significance.

**Figure 3.**
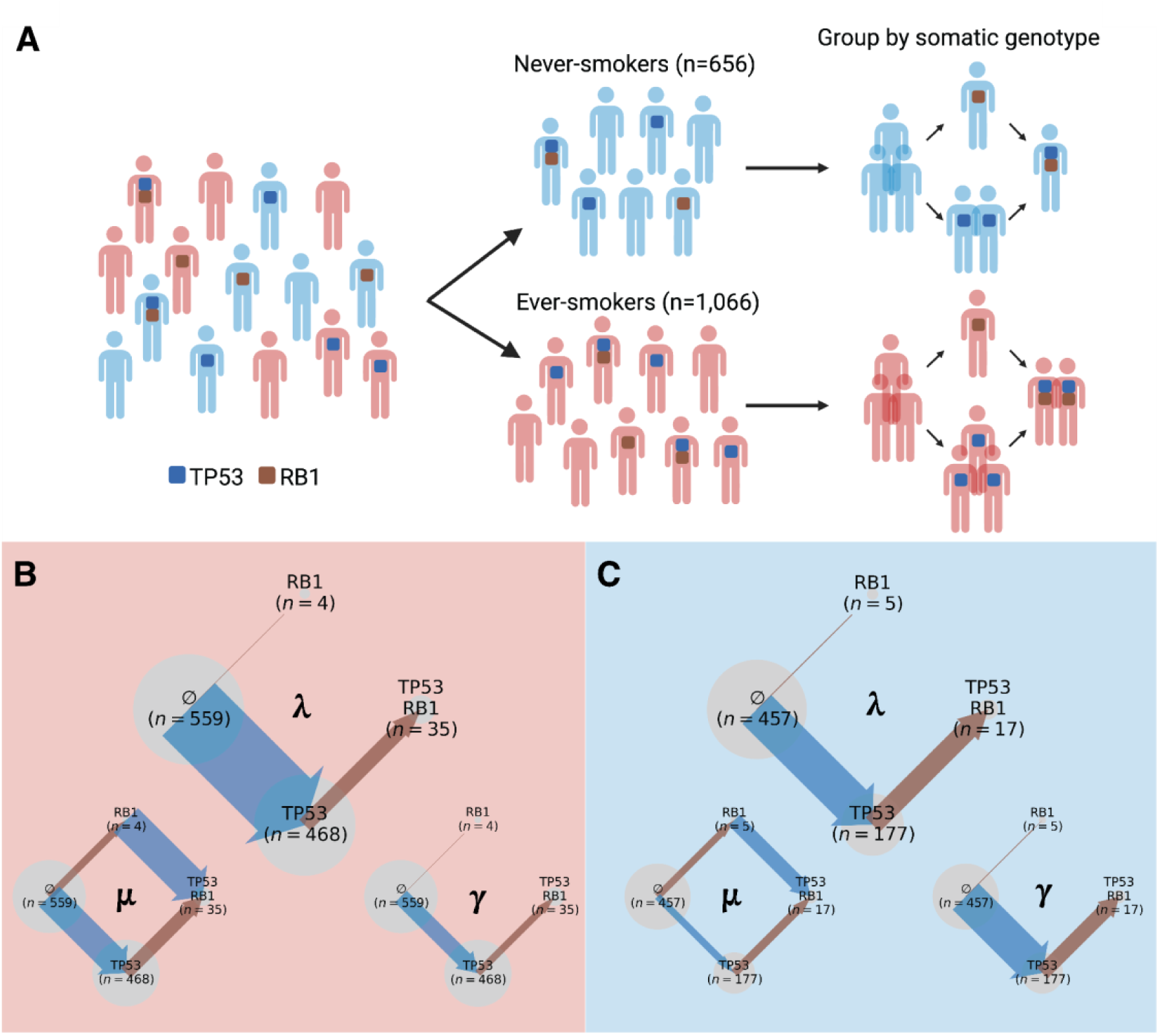
Pairwise selective epistasis inferred within the evolutionary trajectories of lung adenocarcinoma (LUAD), depicted for *TP53* and *RB1*. (**A**) Genomic sequence data from lung adenocarcinomas can be stratified by clinically- and genetically-determined patient smoking status (blue: never-smokers; red: ever-smokers) and somatic genotype. Analysis of (**B**) ever-smoker LUAD and (**C**) never-smoker LUAD deconvolves somatic genotype prevalences (gray circle radius) into genotype-specific fixation rates (λ; arrow width), oncogenic mutation rates (μ; arrow width), and strengths of somatic selection (γ; arrow width) on mutations of *TP53* (dark blue arrows) and *RB1* (brown arrows).

In NS-LUAD, the fixation rate for *TP53* mutations from a wild-type genotype was lower than in ES-LUAD, and the fixation rate for *RB1* mutations in a *TP53*-mutant background was higher (*λ*, **Fig. 3C**). In single-mutant tumors, the mutation rate of *TP53* is nearly double that of *RB1* (*µ*_RB1→RB1+TP53_ *=* 4.0 × 10^−7^, *µ*_TP53→RB1+TP53_ *=* 2.2 × 10^−7^), but in wild-type NS-LUAD tumors, the mutation rate of *TP53* was similar to the mutation rate of *RB1* (*µ*_WT→TP53_ *=* 1.48 × 10^−7^, *µ*_WT→RB1_ *=* 1.8 × 10^−7^). This similarity of mutation rates contrasts with the disparity in ES-LUAD. Despite these differences, the inferred evolutionary trajectory mirrored that of ES-LUAD: the most likely (*λ*) and most adaptive (*γ*) path involved mutation of *TP53* followed by mutation of *RB1*. As in ES-LUAD, prior *TP53* mutation significantly synergistically enhanced selection for *RB1* mutation, and prior *RB1* mutation antagonized selection for *TP53* mutation, without reaching statistical significance (*γ*, **Fig. 3C**).

#### Synergistic epistasis is widespread and often substantial in ES-LUAD

Analysis of all pairwise combinations of the 21 reputed LUAD driver genes revealed frequent synergistic and antagonistic epistasis. In ES-LUAD, synergistic interactions were especially pronounced: for genes whose mutations were under strong positive selection (with the notable exceptions of *TP53* and *KRAS*) synergistic epistasis led to two- to eight-fold increases in selection. For example, selection on mutations in *KEAP1, ATM*, *BRCA2*, *PIK3CA*, and *ARID1A* was markedly enhanced in the presence of prior *TP53* or *KRAS* mutations (**Fig. 4A**). Additional strong synergistic interactions included those affecting mutants of *ARID1A* in the context of somatic genotypes with *ATM* or *RBM10* mutations; mutants of *STK11* in the context of mutated *KEAP1*; and mutants of *ATM* in the context of mutated *STK11*. Mutations in the non-*TP53* tumor suppressors *BRCA2*, *SETD2*, *RBM10*, *RB1*, *MGA*, and *APC*, as well as in the chromatin-remodeling genes *ARID1A* and *SMARCA4*, experienced moderate baseline selection that was frequently enhanced by the presence of diverse prior mutations. These genes never experienced antagonistic epistasis with each other and rarely with other mutations.

**Figure 4.**
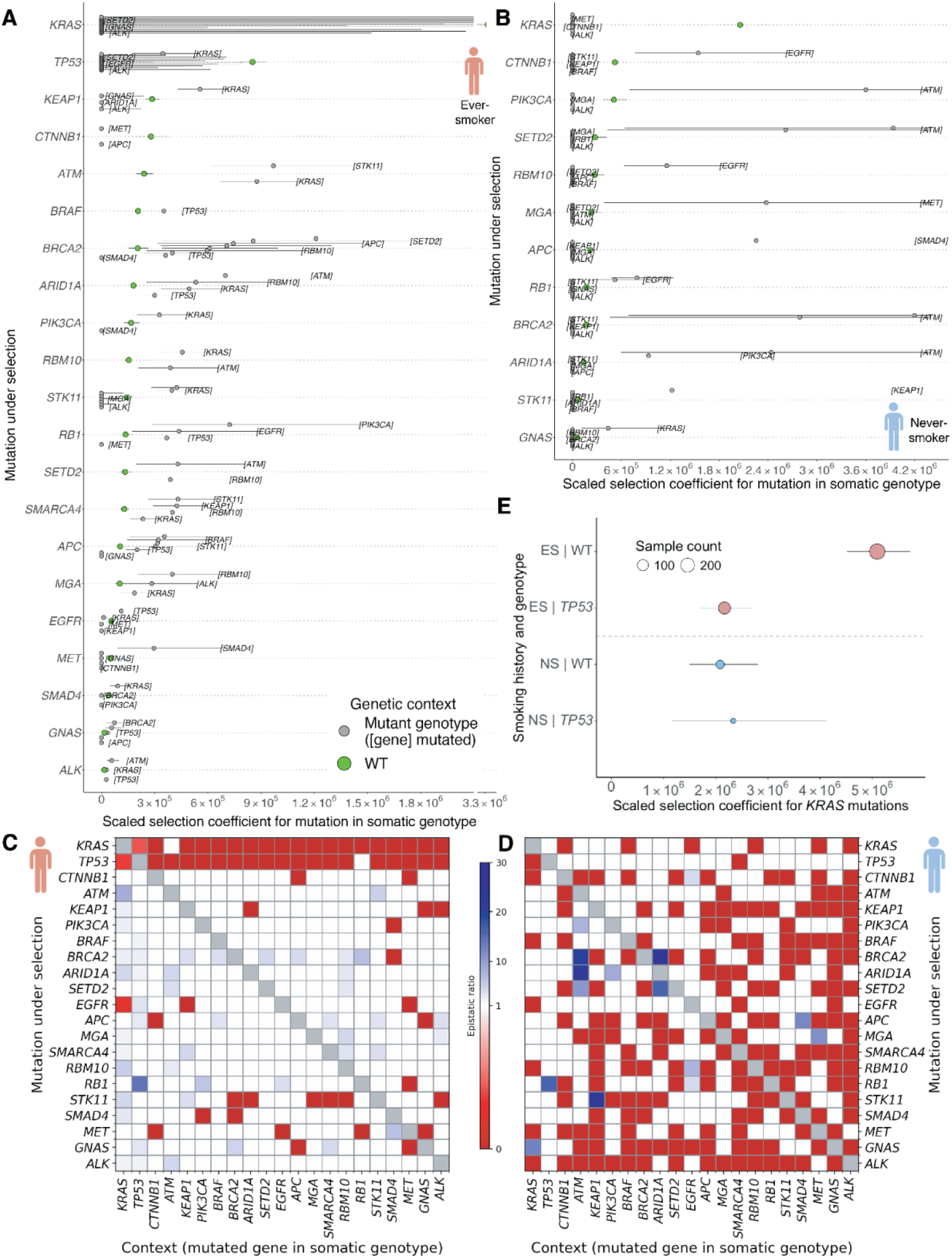
Pairwise epistasis in ever-smoker (ES) and never-smoker (NS) lung adenocarcinoma (LUAD). Strengths of somatic selection for mutations in wild-type tissue (green) and in previously mutated tissue (gray, labeled with previous mutation in brackets) in (**A**) ES-LUAD and (**B**) NS-LUAD. Significant pairwise epistatic interactions depicted in heatmap form for (**C**) ES-LUAD and (**D**) NS-LUAD based on non-overlapping confidence intervals for epistatic strengths of selection. Heatmap values are ratios of selection coefficients for mutations (in each row) from a somatic genotype containing a mutation in another gene (in each column) to the selection coefficient from a somatic genotype lacking the mutation. (**E**) Strengths of somatic selection for mutation of *KRAS* in *TP53*-mutant and *TP53*-wild-type somatic genotypes in ES-LUAD (red) and NS-LUAD (blue).

Patterns of epistasis involving oncogenes were heterogeneous. Notably, *KRAS* and *CTNNB1* mutations showed no elevation of selection in the presence of any prior mutations in the 21-gene set. Conversely, selection for mutations of *GNAS* and *ALK* was enhanced by three prior mutations (**Fig. 4A**). For many oncogenes, selection was enhanced by prior *TP53* mutation, but was antagonized by mutations in other oncogenes such as *MET*. Across all tested genes, the median effect size of statistically significant synergistic epistasis was a 3.6× increase in selection, and the magnitude of this synergy did not correlate with the baseline strength of selection for either the first or the second mutation.

#### Antagonistic effects can reduce positive selection or induce negative selection on drivers

In ES-LUAD, we identified eight gene pairs exhibiting statistically significant antagonistic sign epistasis, in which selection for a driver mutation shifted from positive to negative in the presence of another mutated gene. Mutations in *CTNNB1* and *GNAS* became selectively disadvantageous in the presence of *MET* and *APC* mutations. Similarly, selection for *SMAD4* mutations was antagonized by prior *BRCA2* and *PIK3CA* mutations, and *MET* mutations were antagonized by mutations of *RB1*, *EGFR*, *CTNNB1*, and *GNAS*. In each of these eight cases, the reciprocal mutation order also induced negative selection (e.g., mutated *CTNNB1* induced negative selection for *MET* mutations). For over a hundred other ordered pairs, antagonistic sign epistasis was estimated, without reaching statistical significance. Antagonistic shifts in the magnitude of selection, in which selection for a mutation remained positive but was substantially reduced, were indicated in 33 cases, of which three were significant: the presence of *KRAS* mutations caused a 64% reduction in selection for *TP53* mutations and an 81% reduction in selection for *EGFR* mutations, and *TP53* mutations caused a 58% reduction in selection for *KRAS* mutations (**Fig. 4A**). Across all analyses of ES- and NS-LUAD, new mutations in *TP53* and *KRAS* were never inferred to experience significantly stronger selection in the context of another driver mutation.

#### Antagonistic epistasis is more frequent in NS-LUAD

Compared to ES-LUAD, NS-LUAD exhibited a higher frequency of antagonistic epistasis and fewer instances of synergistic epistasis (**Fig. 4B**). Of the 65 gene pairs exhibiting significant antagonistic epistasis in ES-LUAD, 25 also reached significance in NS-LUAD, and an additional 36 exhibited antagonistic trends without achieving significance (**Fig. 4C–D**). Thus, instances of selective antagonism inferred in ES-LUAD were nearly always present in NS-LUAD.

This consistency contrasted with a substantial inconsistency in selective synergies between ES- and NS-LUAD. Of the 42 gene pairs whose mutations exhibited significant synergistic epistasis in ES-LUAD, only five were significantly synergistic in NS-LUAD. An additional 25 trended synergistic without reaching significance. Strikingly, 16 of these 42 instances were inferred to be significantly antagonistic in NS-LUAD, including many involving *BRCA2*. For example, *KEAP1* mutation enhanced selection for *BRCA2* mutation in ES-LUAD but antagonized it in NS-LUAD (**Fig. 4C–D**). These reversals reveal distinct adaptive constraints imposed by the smoking-altered tumor microenvironment. Overall, antagonistic interactions were relatively stable across smoking status, whereas synergistic epistasis was highly contingent on smoking status.

Mutations in these 21 driver genes were subject to a broad range of epistatic and environmental interactions, as well as combinations thereof (**Fig. S2**). For example, *TP53* mutations antagonized *KRAS* mutations in ES-LUAD but had no substantial effect on *KRAS* mutations in NS-LUAD (**Fig. 4E**). These context-dependent epistatic effects substantially contribute to the divergent trajectories of ES- and NS-LUAD.

Fewer examples of synergistic epistasis were observed in NS-LUAD compared to ES-LUAD. Nevertheless, synergistic epistasis contributed to a median twelve-fold increase in selection on secondary mutations. Notable examples include enhanced selection for *CTNNB1*, *RB1*, and *RBM10* mutations after *EGFR* mutation, for *PIK3CA*, *BRCA2*, *SETD2*, and *ARID1A* mutations after *ATM* mutation, and for *SETD2* mutation after *ARID1A* mutation (**Fig. 4B**). Even in relatively less mutated NS-LUAD, cooperative genetic interactions dramatically reshape the fitness landscape of evolving tumors.

#### Pairwise epistasis can be asymmetric in both sign and magnitude

In both ES- and NS-LUAD, several gene pairs exhibited asymmetric epistasis, in which the direction or strength of interaction depended on the order of mutation acquisition. For instance, in ES-LUAD, mutation of either *KEAP1* or *STK11* enhanced selection on mutations of the other, but with markedly different magnitudes: primary *STK11* mutation was estimated to increase selection for a secondary *KEAP1* mutation by 140%, whereas primary *KEAP1* mutation significantly increased selection for a secondary mutation of *STK11* by 360%. In other cases, the sign of epistasis reversed depending on mutational order. Most notably in ES-LUAD, primary *KRAS* mutation synergistically increased selection for a secondary *KEAP1* mutation, but primary *KEAP1* mutation strongly antagonized selection for a secondary *KRAS* mutation (**Fig. 4A**). This asymmetry means that a mutation in *KRAS* followed by a mutation in *KEAP1* was advantageous, but a mutation in *KRAS* was disadvantageous or lethal when preceded by a mutation in *KEAP1*. This dramatic directional asymmetry and eighteen other such instances exemplify how somatic evolutionary trajectories can be constrained or potentiated by the specific sequence in which mutations occur.

### Higher-order epistasis shapes evolutionary trajectories of ES-and NS-LUAD

#### *KRAS*, *KEAP1*, and *STK11* constitute an epistatically interacting triad in ES-LUAD

Having identified *KRAS*, *KEAP1*, and *STK11* as experiencing strong, ever-smoker-specific selection, we next investigated whether their interactions extended beyond pairwise epistasis to form a higher-order epistatic network. Mutations in all three genes exhibited substantial fixation rates from nearly any initial somatic genotype, particularly en route to the triple-mutant genotype. Fixation rates for *KRAS* mutations were estimated to be very low when arising after *KEAP1* or *STK11* mutations alone compared to when arising as the initial mutation or in the *KEAP1*-*STK11* double-mutant (**Fig. 5A**). Nevertheless, these three genes form an epistatic triad in which mutations in any one tend to increase selection for the others.

**Figure 5.**
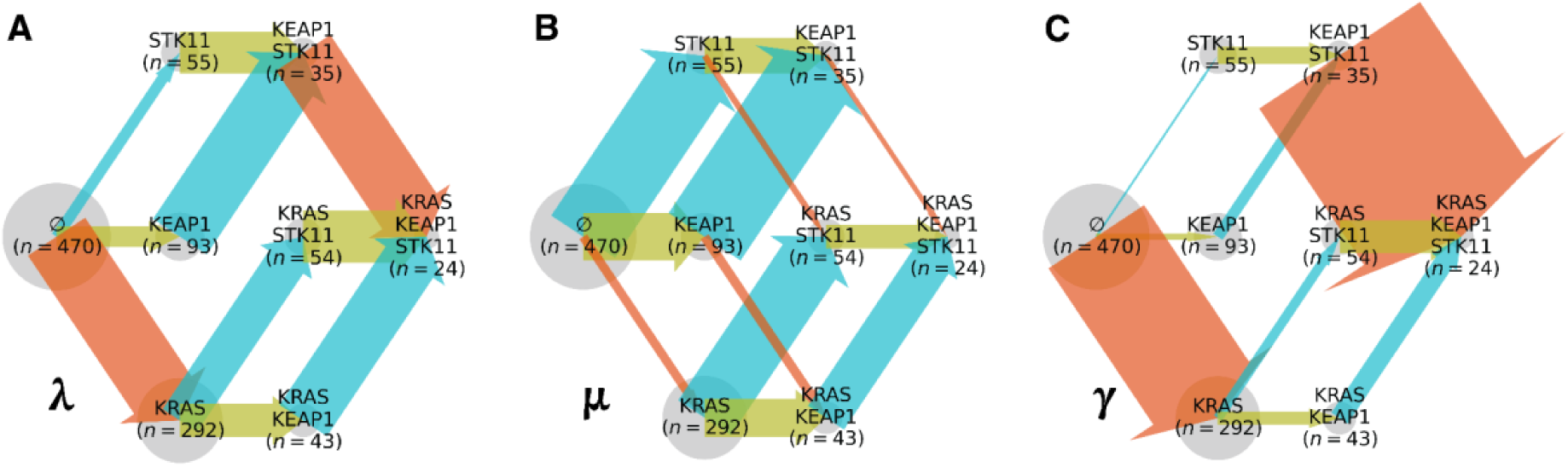
Evolutionary trajectory for *KRAS*, *KEAP1*, and *STK11* mutations in ever-smoker lung adenocarcinoma. (**A**) Fixation rates, (**B**) oncogenic mutation rates, and (**C**) scaled selection coefficients for mutations of *KRAS* (orange arrows; arrows width proportional to rate or coefficient), *KEAP1* (light blue arrows), and *STK11* (yellow arrows) conditioned on somatic genotype (gray circles; radius proportional to genotype prevalence).

Among the three genes, *STK11* manifested the highest oncogenic mutation rate at each progressive step, followed by *KEAP1*, and then *KRAS* (**Fig. 5B**). Given these differential underlying rates of mutation, we computed the consequent scaled selection coefficients for mutations in each genotype in the three-gene model, and compared selection between genotypes to quantify pairwise and higher-order selective epistasis. Many tumors possessed mutations in at least one of these genes (**Fig. 5C**, second column of nodes), providing high power to detect pairwise epistasis. Indeed, our three-gene model recapitulated the pairwise epistatic relationships inferred from the two-gene models (**Fig. 4C**). However, concurrent mutations of two or more of these genes were relatively infrequent (**Fig. 5C**, third and fourth columns of nodes), limiting power to detect higher-order epistasis with statistical confidence. For example, substantial uncertainty in the selection estimate for *KRAS* mutations in tumors harboring *STK11* and *KEAP1* mutations precluded a definitive inference of higher-order interactions. In contrast, because of the sufficiently high prevalence of *KRAS*-*KEAP1*- and *KRAS*-*KEAP1*-*STK11*-mutant genotypes, we were able to infer higher-order epistatic effects on *STK11* with greater confidence.

*STK11* mutations experienced significantly stronger selection in the presence of either *KRAS* or *KEAP1* mutations. If the synergistic effects of *KRAS* and *KEAP1* mutations on *STK11* compounded additively, then *STK11* mutations would experience selection equivalent to the three-way product of the individual synergistic effects and the baseline selection for *STK11* mutations. However, the upper bound for selection for *STK11* mutations in a *KRAS*-*KEAP1* double-mutant genotype was less than this product (**Fig. 5C**). Accordingly, the compound synergistic effect of concurrent *KEAP1* and *KRAS* mutations represents an example of sub-additive higher-order epistasis.

#### Most pairwise epistatic effects compound without strong higher-order interaction

The sub-additive epistasis affecting mutations of *STK11* demonstrates that assessments of higher-order epistasis will be necessary to thoroughly understand cancer evolution. However, most effects of driver mutations compounded in a manner consistent with minimal higher-order epistasis. Examining all 1330 triads out of the 21 genes, we compared selection for a new mutation in single-versus double-mutant backgrounds. In the vast majority of cases, a combination of pairwise epistatic effects was sufficient to explain significant differences in selection for a new mutation between single- and double-mutant genotypes. Generally, when selection for a secondary mutation in one gene was epistatically altered by presence of a primary mutation in a second gene and was unaffected by a primary mutation in a third gene, selection on the secondary mutation remained commensurately epistatically altered when both other variants of the triad were concurrently present. For example, in ES-LUAD, selection for *SMARCA4* mutations was synergistically enhanced by *STK11* mutations but unaffected by *TP53* mutations, and correspondingly, selection for mutations in *SMARCA4* was synergistically increased in a *STK11*-*TP53*-mutant genotype (**Fig. 6A,B**). Similarly, in NS-LUAD, selection for *RBM10* mutations was synergistically increased in both a *TP53*-mutant and *TP53*-*EGFR*-mutant genotype (**Fig. 6C**). Overall, pairwise interactions typically dominate the adaptive landscape of LUAD, but higher-order epistasis can have substantial influence on specific evolutionary trajectories.

**Figure 6.**
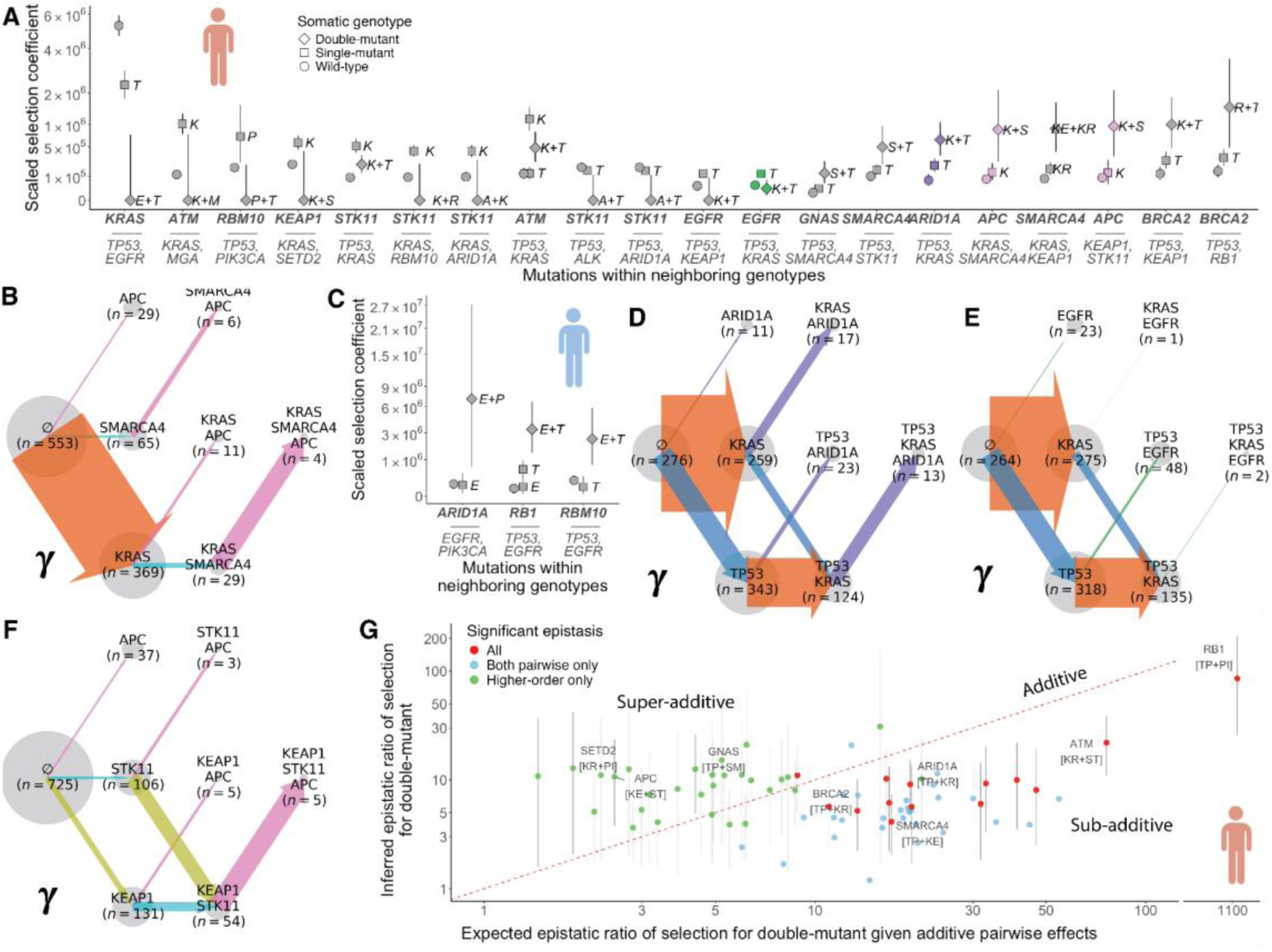
Higher-order selective epistasis in lung adenocarcinoma (LUAD). (**A**) Selection within gene triads (*y* axis is square-root scaled), for a mutation (bold font) in a context of no co-occurring mutations (circle), one co-occurring mutation (square; labeled with first letter(s) of the gene(s) indicated in the tick label that are mutated), and two co-occurring mutations (diamond) in ever-smoker (ES) LUAD [pink: visualized in panels B and F; purple: visualized in panel D; green: visualized in panel E]. (**B**) In ES-LUAD, the synergistic effect of *SMARCA4* mutations (light blue arrows) on selection for *APC* mutations (pink arrows; arrow width proportional to scaled selection coefficient; circle radius proportional to somatic genotype prevalence) does not diminish in the presence of a neutrally-interacting *KRAS* mutation (orange arrows). (**C**) Selection within gene triads in never-smoker (NS) LUAD; *y* axis is square-root scaled, symbols are as in panel A]. (**D**) Pairwise and higher-order epistasis in ES-LUAD: co-mutation of *TP53* (blue) and *KRAS* (orange), each of which independently increase selection for *ARID1A* mutations (purple), produces an even stronger synergy for *ARID1A* mutations; and (**E**) antagonistic effect of *KRAS* mutations (orange) on selection for *EGFR* mutations (green) outweighs the synergistic effect of *TP53* mutations (blue). (**F**) In ES-LUAD, co-mutation of *KEAP1* (light blue) and *STK11* (yellow) synergistically increases selection for *APC* mutations (pink) despite neutral pairwise effects. (**G**) Comparison of the epistatic effect of double-mutant-genotype (label: bottom row, two-letter abbreviation) on selection for new mutation (label: top row) to the “expected” product of pairwise epistatic effects (blue points: both pairwise epistatic effects are statistically significant, but not the higher order effect; green: the higher-order effect is significant, but neither pairwise effect is; red: all effects are significant)

When two co-occurring mutations each independently enhanced selection for a third mutation, the synergistic effects frequently compounded. For instance, *ARID1A* mutations experienced increased selection in both KRAS- and *TP53*-mutant genotype. These synergistic effects compounded, resulting in mutations of *ARID1A* experiencing three times stronger selection in a *KRAS*-*TP53*-mutant genotype than in a *TP53*-mutant genotype and nine times stronger selection than in a wild-type genotype (**Fig. 6D**). Similar patterns of compounded synergy were evident for *ATM* mutations in *KRAS*-*STK11*-mutant genotypes, as well as for *SMARCA4* mutations in a *KEAP1*-*STK11*-mutant background. Across cases, compounded synergy frequently drove selection to levels up to an order of magnitude higher than in genotypes lacking either partner mutation. This substantial compounding constitutes a crucial element of how driver mutations accumulate in cancer genomes, positively reinforcing progression of cancer toward more aggressive forms.

#### Higher-order epistasis often manifests as sub-additive or emergent synergistic interactions

In the case of compounding synergistic effects, it is also possible to observe super-additive higher-order epistasis, in which selection for a mutation in the presence of two independently synergistic mutations is even larger than what would be expected from the product of their independent synergistic effects. Indeed, some cases are consistent with either additive synergy or super-additive epistasis, such as mutations of *SMARCA4* in a *KRAS*-*KEAP1*-mutant genotype in ES-LUAD (**Fig. 6A**). More frequently, cases are consistent with either additive synergy or sub-additive epistasis, in which selection for a mutation in the presence of two independently synergistic mutations is smaller than expected from the sum of synergistic effects. Indeed, in 43 of 45 cases wherein a new mutation arose in a genotype of two significantly synergistic mutations, estimates of selection for the new mutation were lower than would be expected from purely additive synergistic epistasis. Such widespread sub-additivity pervades the adaptive landscape, tempering rather than amplifying selective pressures, even in the context of strong pairwise synergy.

We also identified cases of higher-order epistasis involving the co-occurrence of synergistic and antagonistic partner mutations, as well as the emergence of synergy in the co-occurrence of neutrally-interacting partner mutations. In ES-LUAD, for example, selection for *EGFR* mutations was synergistically enhanced by prior *TP53* mutations but antagonized by *KRAS* mutations. In a *TP53*-*KRAS*-mutant genotype, *EGFR* mutations experienced significantly less selection than in a *TP53*-mutant *KRAS*-wildtype genotype (**Fig. 6A**,**E**). In fact, these *EGFR* mutations experienced levels of selection similar to those they would experience in a genotype lacking either mutation, indicating a neutralizing result of the combination of synergistic and antagonistic epistasis.

Another form of higher-order epistasis is the emergence of synergy or antagonism from a combination of mutations, each of which does not individually exhibit pairwise epistasis with the affected gene. We observed several instances in which selection for a mutation was significantly increased by the co-occurrence of two mutations that did not individually have significant synergistic effects: in ES-LUAD, selection for mutations of *APC* increased by an order of magnitude in a *KEAP1*-*STK11*-mutant genotype despite not being significantly altered when only one of *KEAP1* or *STK11* was mutated (**Fig. 6A**,**F**). In this instance, significance was reached for synergy when compared to wild-type or a *KEAP1*-mutant genotype, but not when compared to the *STK11*-mutant genotypes, which was itself epistatically neutral. We detected multiple similar instances of emergent synergy, such as for *ARID1A* mutation in an *EGFR*-*PIK3CA*-mutant genotype in NS-LUAD (**Fig. 6G**). We also observed one instance of emergent antagonism, in which *STK11* mutations were antagonized in a *TP53*-*EGFR*-mutant genotype but not in either single-mutant genotype alone.

Overall, we observed two major classes of higher-order epistasis: sub-additive synergy and emergent—and consequently super-additive—synergy. In particular, when both component epistatic effects were strongly synergistic, their combination was almost always sub-additively synergistic (**Fig. 6G**). On the other hand, when both component epistatic effects were neutral or insignificant, a super-additive synergistic effect occasionally emerged from their combination (**Fig. 6G**). Interestingly, as the strength of the component epistatic effects increases, the inferred epistatic effect of the double-mutant combination rapidly plateaus at an approximately ten to twenty-fold increase in selection.

In this survey of the adaptive landscape of LUAD, we discovered several cases of higher-order epistasis that substantially influence the selective advantage of oncogenic mutations. Over a third of LUAD samples in ever-smokers possessed mutations in three or more of 21 tested genes—a proportion that would likely rise in an analysis of a greater number of driver genes. In these patients, higher-order epistasis may play a substantial role in the evolutionary trajectories of their tumors and their responses to targeted therapies. Indeed, we found that a wide variety of adaptive trajectories ensued from the combined epistatic effects of multiple mutations. Some of these trajectories cannot be predicted from pairwise analyses. However, our large-scale analysis of the adaptive landscape of 21 mutations has significantly supported only a subset of the many potential instances of higher-order epistasis between oncogenic mutations in LUAD. Indeed, multiple other instances of higher-order epistasis were estimated to be present. Their effect would be clarified with greater sample sizes.

## Discussion

Here we have characterized the distinct adaptive landscapes of lung adenocarcinoma in ever-smokers and never-smokers by first estimating gene-specific background mutation rates in ES- and NS-LUAD, then comparing the intensity of somatic selection and selective epistasis acting on those mutations between the two groups. Consistent with the established mutagenic effects of tobacco carcinogens [4], we detected elevated oncogenic mutation rates in all tested genes in ES-compared to NS-LUAD. Accounting for these background rates revealed that mutations in genes such as *EGFR*, *PIK3CA*, and *MET* provide a stronger selective advantage in NS-LUAD, while mutations in *KRAS*, *KEAP1*, and *STK11* provide a stronger selective advantage in ES-LUAD. Focusing on *EGFR* and *KEAP1*, we found that these adaptive differences were at least partially explained by the differential effects of these mutations on epithelial-mesenchymal transition and DNA replication transcriptional program activity in ES-and NS-LUAD.

By quantifying the selective advantage of these mutations within specific somatic genotypes, our analysis recovered known epistatic interactions—such as the enhancement of selection on *RB1* and *EGFR* mutations by prior *TP53* mutations—and revealed context-specific interactions, such as the synergistic effect of *EGFR* mutations on *CTNNB1* mutations uniquely in NS-LUAD. We also detected substantial asymmetries: *CTNNB1* mutations did not impact *EGFR* mutations in ES- or NS-LUAD. Notably, we found more frequent synergy in ES-LUAD and substantially more pervasive antagonism in NS-LUAD, constituting fundamental differences in the evolutionary navigability of their adaptive landscapes. Finally, we showed that higher-order epistasis—particularly sub-additive synergistic and emergent synergistic epistasis—is frequent in and substantially guides the evolutionary trajectories of cancer. These findings of differential selection and epistasis reveal that exposure to tobacco smoke reshapes the adaptive landscape of LUAD not only by elevating mutation burden, but also by altering the selective value of mutations and their epistatic interactions—highlighting opportunities to leverage these adaptive differences for biomarker development and therapeutic stratification.

The prevalences of driver mutations have been reported to substantially differ between smokers and never-smokers in LUAD [21,22] as well as in non-small cell lung cancer [NSCLC; 23,24]. However, previous studies have either refrained from explaining these differences or attributed them primarily to tobacco-induced mutagenesis [22,23]. Our results show that they often cannot be explained by differential mutagenesis alone. For example, *KRAS* variants are over five times more frequent in ES- than NS-LUAD tumors, but the background, neutral rate at which *KRAS* mutations occur is only about three times higher in ES-LUAD. To resolve this discrepancy, *KRAS* mutations must be nearly twice as oncogenic in ES-compared to NS-LUAD. In general, the differential prevalence of these driver mutations is often not solely attributable to tobacco-induced mutagenesis, but also due to their altered selective advantage in the tobacco-altered lung environment. This physiological-adaptive effect of smoking is crucial to account for: changes in the selective advantage of a mutation suggest altered functional impact within a given tissue environment, with direct implications for the efficacy of targeted therapies and the interpretation of biomarkers in smokers versus non-smokers.

Multiple perspectives have proposed that the adaptive landscape of oncogenesis is altered by physiological insults from endogenous and exogenous factors [14,17,52,53]. However, data-driven evidence for this hypothesis has remained limited [cf. 54]. Here, we provide compelling evidence that canonical driver mutations confer altered selective advantages in the smoking-affected lung microenvironment of ever-smokers with LUAD. These shifts in selection are linked to context-dependent phenotypic effects of mutations. For example, *EGFR* mutations are associated with greater activity of the epithelial-mesenchymal transition (EMT) transcriptional program in NS-LUAD, but not in ES-LUAD—likely explaining their stronger selective advantage in never-smokers. Such divergent phenotypic consequences can be attributed to smoking-induced alterations to transcriptional program activity [55–57], tumor microenvironment composition [58], and cell fitness. These findings provide a mechanistic basis for how exogenous exposures can reshape the evolutionary trajectory of cancer by altering the adaptive value of oncogenic mutations.

In the case of *TP53*, Rodin and Rodin [13] reported comparable background mutation rates between smokers and nonsmokers with lung cancer, and attributed the higher frequency of *TP53* mutations in smokers to increased selection in the smoking-altered microenvironment. By contrast, our analysis—leveraging a substantially larger dataset, incorporating tissue- and context-specific covariates, and applying advanced statistical methodologies to estimate background mutation rates [6]—identified a markedly elevated *TP53* mutation rate in ES-LUAD. With this substantial differential mutagenesis accounted for, our analysis revealed that *TP53* mutations are actually under weaker positive selection in ever-smokers than in never-smokers. This finding challenges earlier conclusions and underscores the importance of accurately resolving both mutational processes and selection when interpreting variant prevalence across environmental exposures.

Despite mounting evidence of the substantial biological impact of tobacco smoke, clinical trials for lung cancer therapies often do not assess the influence of smoking status—and frequently do not even collect usage history [59], perhaps due to a conception that tobacco smoke has a negligible effect on outcomes [60]. However, our findings reveal that tobacco smoke profoundly reshapes the evolutionary trajectory of lung adenocarcinoma, necessitating more systematic incorporation of smoking history into trial design and treatment decision-making, particularly for targeted therapies. For example, the substantially stronger selection for *EGFR* mutations in NS-LUAD implies that EGFR tyrosine kinase inhibitors (TKIs) should yield greater clinical benefit in never-smokers. Indeed, a retrospective analysis of patients with *EGFR*-mutant LUAD found that the objective response rate (ORR) of patients receiving EGFR TKIs declined from 73% among never-smokers to 46% among individuals with over 10 pack-years of smoking history [61]. Similar patterns have been reported in additional retrospective studies of LUAD [62] and across NSCLC more broadly [63,64]. This concordance between selection-based predictions and observed outcomes highlights the translational value of selection coefficients as predictors of targeted therapy efficacy. By identifying likely responders based on environmental exposures that shape the tumor microenvironment, exposure-informed selection coefficients can improve clinical trial stratification, enhance statistical power, and refine treatment personalization in precision oncology.

Prior studies have questioned whether computational analyses can reliably detect epistasis, citing concerns about confounding by subtype-specific mutation profiles, variation in mutation burden, and limited statistical power [31,65]. To address these concerns, we analyzed ES- and NS-LUAD independently and comparatively, modeled subtype- and gene-specific background mutation rates and effect sizes of mutations in specific somatic genotypes instead of suboptimal measures of co-occurrence or mutual exclusivity, and integrated data from multiple cohorts to ensure sufficient statistical power. Consequently, we were able to make robust inferences of both pairwise and higher-order selective epistasis, many of which align with experimental and clinical evidence. For example, consistent with our finding that *KRAS* mutations synergistically enhance selection for *KEAP1* and *STK11* mutations, prior studies show that *KEAP1* loss accelerates LUAD progression in *KRAS*-mutant mouse tumors [66,67] and compensates for the deleterious effects of *KRAS*-*STK11*-co-mutation [68], and mutations of *KEAP1* and/or *STK11* worsen survival and treatment response in *KRAS*-mutant LUAD [69,70] and NSCLC [71–76]. Furthermore, in accordance with the inferred mutually synergistic effects of mutations of *KEAP1* and *STK11* in ES-LUAD, co-mutation of *KEAP1* and *STK11* promote cell proliferation [77] and worsen survival in NSCLC independently of *KRAS* [78,79].

Our analyses also identified synergies between *EGFR* and other drivers: *RBM10* and *CTNNB1* in NS-LUAD, and *RB1* in both NS- and ES-LUAD. These interactions are corroborated by experimental and clinical studies showing that *RBM10* mutations promote *EGFR*-mutant LUAD [80] and confer resistance to EGFR TKIs [81], that *CTNNB1* mutations are enriched in advanced *EGFR*-mutant NSCLC and associated with metastasis and TKI resistance [82], and that *RB1* mutations in *EGFR*-mutant LUAD mediate histologic transformation to small-cell lung cancer [83,84]. We also revealed that mutations of *TP53* and *KRAS* were widely antagonized by other driver mutations, consistent with their early, clonal emergence during LUAD evolution [85,86] and supporting their frequently primary role in tumor initiation. This panoply of alignments between our results and experimental and clinical evidence encourages deeper investigation into the many epistatic relationships we have revealed that remain untested in the laboratory or clinic. They represent fertile ground for mechanistic investigation and promise new therapeutic targets and biomarker candidates.

Previous studies have shown that epistatic interaction networks vary across cancer types in humans [87] and across environmental contexts in other organisms [88–93]. However, extensive differences in selective epistasis within a single cancer type exposed to distinct tissue environments have not previously been demonstrated. Here, we have shown that tobacco smoke exposure markedly reshapes the selective epistatic interactions underlying LUAD evolution. Broadly, synergistic epistasis was more frequent in ES-LUAD, whereas antagonistic epistasis was more frequent in NS-LUAD. Indeed, in some cases, epistasis between driver genes was synergistic in ES-LUAD, but antagonistic in NS-LUAD. In other cases, the magnitude of synergy differed across exposures: synergy for mutation of *TP53* in *EGFR*-mutant tumors was stronger in ES- than in NS-LUAD. The higher frequency of synergy and lower frequency of antagonism in ES-LUAD suggests a more navigable adaptive landscape—characterized by a higher frequency of beneficial mutational combinations and fewer evolutionary barriers—than in NS-LUAD. This increased accessibility enables ES-LUADs to develop along a greater diversity of evolutionary trajectories, contributing to increased interpatient heterogeneity. Moreover, it may also help to explain why tobacco smoke elevates the risk of lung cancer to such an exceptional degree. In NS-LUAD, widespread antagonistic epistasis may reflect a greater fitness cost of mutation accumulation, possibly due to a more immunologically reactive tumor microenvironment [12,48,58,94] or due to a lower tolerance for mutation accumulation in the non-dysregulated lung tissue cells of never-smokers. Taken together, these findings reveal that environmental exposures modulate selective epistasis within a cancer type, suggesting the importance of exposure history to predicting tumor evolutionary trajectories and to guiding epistasis-informed treatment strategies.

It has often been assumed that the adaptive landscape of cancer can be adequately described without epistasis [6,95–99] or with only pairwise epistasis [31,32,100]. However, our results demonstrate that not only pairwise but also higher-order epistasis—particularly sub-additive and emergent epistasis—play a crucial role in modulating the adaptive value of cancer drivers. For instance, in ES-LUAD, selection for *APC* mutations increased by an order of magnitude in an *KEAP1*-*STK11*-mutant genotype, despite showing no significant pairwise synergy with either mutation alone. This emergent epistasis, arising only in the triadic context, would be missed by analyses restricted to pairwise comparisons, necessitating consideration of multi-mutant genotypes in both computational and experimental studies. Moreover, almost all significant pairwise synergistic effects combined sub-additively in triads, rapidly reaching a plateau of synergistic benefit. This pattern aligns with the diminishing-returns epistasis observed in asexual populations [44–47]. Overall, higher-order synergistic epistasis biases tumors toward some evolutionary trajectories, while higher-order antagonism prunes most trajectories. Both make tumor evolution more predictable. Thus, incorporation of higher-order epistasis into models of tumorigenesis is essential to understanding and guiding the evolutionary paths that tumors take under therapy.

Current NSCLC treatment guidelines for EGFR TKIs do not account for co-occurring mutations. However, our findings suggest that concurrent mutations could serve as biomarkers of treatment response. Antagonistic epistasis between *EGFR* mutations and mutations in *KRAS* or *KEAP1* implies that the efficacy of EGFR TKIs may be reduced in tumors harboring these concurrent alterations. This prediction aligns with clinical observations: *KRAS* mutations decrease survival in patients with *EGFR*-mutant LUAD patients treated with EGFR TKIs [101], and *KEAP1* mutations reduced EGFR TKI efficacy in *EGFR*-mutant lung cancer [102,103]. More broadly, antagonistic epistasis is a signal of synthetic lethality [32]. Our results also demonstrate mutual negative selection between *BRCA2* and *SMAD4* loss-of-function mutations—suggesting that pharmacological inhibition of *SMAD4* could be therapeutically beneficial in *BRCA2*-mutant tumors, in analogy to usage of PARP inhibitors in *BRCA1*/*2* deficient tumors. In general, selective epistasis can clarify the frequent heterogeneity of response to targeted therapies [37,104], and can improve stratification of responders and nonresponders to targeted therapy in clinical trials [105,106], enhancing trial power and therapeutic precision.

Because *TP53* mutations synergize with mutations of *EGFR*, we would expect that in tumors with concurrent *EGFR* and *TP53* mutations, EGFR TKIs would be especially effective because they would both eliminate the baseline effect of the *EGFR* mutation and diminish the effect of the concurrent *TP53* mutation that is mediated through the *EGFR* mutation. However, *TP53* mutations are consistently associated with resistance to EGFR TKIs in *EGFR*-mutant LUAD [107–109] and NSCLC [110–114]. This contradiction may be explained by the pleiotropic effects of *TP53* loss, including increased somatic nucleotide mutation rates [109], widespread copy-number alterations [107,115], and dysregulation of gene expression programs linked to resistance [116] and progression [115]. These pleiotropic effects provide greater genetic variation that facilitates the evolution of resistance independently of the selective synergy between *TP53* loss and *EGFR* mutation.

Previous computational studies of pairwise interactions have demonstrated asymmetries in the magnitude of pairwise epistasis [cf. 30]. Our analysis has revealed not only epistasis of asymmetric magnitudes, but also asymmetric signs of selection, in which one ordering of the mutations is advantageous while the reverse is disadvantageous. These directional asymmetries likely arise from cellular or stable transcriptional transitions initiated by the primary mutation that unidirectionally alter the selective context for subsequent alterations—an effect not captured by conventional analyses of mutual exclusivity or co-occurrence. Indeed, the presence of such asymmetries of epistasis are evident in established genetic models of tumor evolution, such as the *APC* → *KRAS* → *TP53* mutation sequence in colorectal cancer [117,118], whose strongly preferential ordering requires directional epistasis. Asymmetric epistasis has also been supported by genetic and computational genomic studies of *TP53* and *CCNE1* mutations [32], *TP53* and *RB1* mutations [30], and *JAK2* and *TET2* mutations [119]. The presence of directional epistasis implies preferential orders of mutation acquisition that offer new opportunities to predict and potentially redirect cancer evolution.

This study was subject to several limitations. First, its retrospective nature introduces the possibility that covariates of smoking behavior—such as comorbid health states, environmental exposures, inherited genetics, and sex biases—may contribute to the observed differences in the adaptive landscapes of ES-LUAD and NS-LUAD. However, none of these factors are known to exert a physiological impact on the lung that is comparable to smoking tobacco. Accordingly, the selective differences we report are most parsimoniously attributed to the direct and downstream effects of smoking. Second, smoking status in our study was primarily inferred from mutational signatures in tumor sequences, which can be susceptible to misclassification. However, this method has been validated for classification of smoking status in NSCLC and is believed to outperform self-reported smoking histories [21,25,120]; self-reported status has been found to be unreliable [121,122] and often does not account for the risk conferred by secondhand smoke [123]. Furthermore, because our approach explicitly quantifies the smoking-associated mutational burden in the lung periphery—where LUAD develops—it also implicitly identifies patients who have experienced significant injury from smoking in the regions where LUAD originates. Finally, our analysis was restricted to somatic SNVs and did not consider indels, copy number alterations (CNAs), or other structural variants (SVs). CNAs are common in cancer [124] and pairwise selective epistasis has been indicated between SNVs and CNAs and other SVs [125,126] or between SVs [37,87,127]. Unfortunately, accurate background mutation rates are currently unavailable for SVs. As such rates become calculable, integrated analysis of SNVs and SVs will reveal new epistatic relationships and clarify the somatic adaptive landscape of cancer.

Overall, we have demonstrated that the adaptive landscapes of ES- and NS-LUAD diverge substantially, implicating tobacco smoking not only as a mutagen but also as a modulator of somatic selection. Our analysis also shows that symmetric and asymmetric selective epistasis among driver mutations is both substantial and pervasive in lung adenocarcinoma and generates the distinct evolutionary trajectories of ES-LUAD and NS-LUAD. Furthermore, we provide a systematic characterization of higher-order selective epistasis in LUAD, revealing its underrecognized role in shaping tumor evolution. Together, these results show that the contribution of mutations to tumor growth is profoundly context-dependent—modulated by both tumor somatic genetics and the environmental exposures it has experienced. This insight illuminates new opportunities for precision oncology through the integration of somatic genotype and exposure history into treatment strategies. Continued investigation into epistatic, environmental, and even gene-by-gene-by-environment interactions in lung adenocarcinoma and other cancers will be essential for the advancement of our understanding of cancer evolution and the improvement of patient-specific therapeutic strategies.

## Methods

### Design overview

The aim of this study was to test for differences in background mutation rates, subsequent somatic selection, and gene-by-gene epistatic effects between ES- and NS-LUAD. To do so, we aggregated mutation data of 2,097 LUAD tumor samples from multiple institutional sources. We classified tumors as either ES- or NS-LUAD either by detection of the smoking-associated mutational signature SBS4 in exome or genome sequences or, in the case of targeted panel sequence data, by relying on the patient’s self-reported smoking history. We applied established approaches to estimate background mutation rates, scaled selection coefficients, and pairwise and higher-order selective epistasis within the ES- and NS-LUAD cohorts. To assess the differential transcriptional effects of driver mutations, we compared ES- and NS-LUAD tumors with and without specific mutations. Differences in selection were conservatively evaluated based on non-overlapping 95% confidence intervals.

### Data sources

We sourced clinicogenomic data from (i) eight published datasets hosted by cBioPortal [85,128–135], (ii) The Cancer Genome Atlas dataset hosted on Genomic Data Commons [136], and (iii) 108 samples sequenced at Yale University [137]. All datasets reported sample sizes, sequencing methods, and clinical covariates, and were incorporated into our analysis accordingly (**Supplementary Table 1**). We only included samples from primary LUAD tumors, and we did not include any datasets from studies that preselected patients based on their mutational profiles. When multiple samples from a single patient were available, one was selected at random. In cases where patients appeared in more than one dataset, we retained the sample with the highest-quality sequencing and most complete clinical annotations.

For consistency across datasets, we standardized all variant genomic coordinates to the hg19 reference genome using the liftOver function of rtracklayer v1.62.0 [138]. We used cancereffectsizeR v2.10.1 to remove variants that failed liftover, were incorrectly classified as single-nucleotide variants (SNVs), or did not meet quality criteria for downstream analysis. For panel-sequenced samples, we inferred the genomic regions covered by each panel from the gene list provided for each panel using the corresponding hg19 coordinates.

### Mutational signature attributions and smoking-status classification

We performed mutational signature deconvolution for tumor samples with whole-exome or whole-genome sequence data, but not for panel-sequenced samples, due to the low reliability of signature attribution when few silent somatic variants are present. We conducted signature attribution using cancereffectsizeR with MutationalPatterns v3.12.0 [139], applying COSMIC v3.2 single-base substitution signatures [26,140]. We excluded COSMIC signatures that were unlikely to have been observed in lung adenocarcinomas [140], and we excluded treatment-associated signatures when deconvolving samples known to be treatment-naive. Samples in which the smoking-associated SBS4 signature contributed more than the MutationalPatterns default threshold of 5% of the total signature burden were classified as ever-smoker ES-LUAD, and otherwise were classified as never-smoker NS-LUAD. Zhang et al. detected no signature of SBS4 in the samples used in their study; therefore we directly classified these samples as NS-LUAD. As in Rosenthal et al. [141], we excluded samples with ≤50 single-nucleotide variants in whole-exome data from classification due to insufficient mutation burden for reliable signature deconvolution. We compared signature-based smoking assignments to available clinical annotation of smoking status across datasets, validating that there was 67–90% concordance. Discrepancies were likely due to underreporting of smoking, second-hand smoke exposure, minimal mutational consequences in light smokers, or misclassification in clinical records. The selection and epistasis results produced from either classification method were 99.4% consistent with each other. We classified panel-sequenced samples according to clinical annotations of smoking status when available; we excluded samples annotated as exposed to “former light” smoking from all smoking-stratified analyses due to ambiguity in exposure and effect.

### Estimation of oncogenic mutation rates

We estimated gene-specific oncogenic mutation rates separately for the ES- and NS-LUAD cohorts following the approach of Alfaro-Murillo and Townsend [28]. The oncogenic mutation rate for a gene was defined as the neutral rate at which oncogenic variants occur within the gene. We defined oncogenic variants as variants that were observed at least once in any tumor sample in the dataset. The background oncogenic mutation rate of each gene was computed as the sum of the background rates of all oncogenic variants within the gene. For a specific variant derived from a trinucleotide mutation type *m* (*e.g.* CCT → CAT) at a site *i* within gene *g* in a tumor sample cohort *c*, the somatic background mutation rate

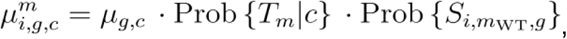

where *μ_g,c_* is the background mutation rate of the entire gene *g* (the rate at which variants occur in *g*) within cohort *c*, *T*_*m*_ is the event that an arising variant is of trinucleotide mutation type *m*, and 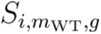 is the event that the variant occurs at site *i* among all other trinucleotide sequences that match the wild-type (WT) allele of *m* (*m*_*WT*_, *e.g.* CCT), within *g*.

The probability of the variant being of type *m* given a cohort *c*, Prob{*T_m_*|*c*}, is computed as the proportional frequency of *m* within cohort *c*, equivalent to the ratio of the number of variants of type *m* in cohort 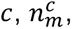 to the total number of variants in *c*, *N*_c_. Assuming an equal probability of such a variant occurring at all possible sites within *g*, the probability of such a variant occurring at site 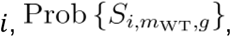 is the inverse of the number of sites within *g* with the same trinucleotide sequence as 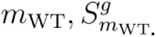 Therefore

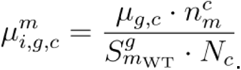

The background oncogenic mutation rate of gene *g* is the sum of the background rates of all oncogenic variants within *g*:

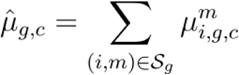

where *S_g_* denotes the set of pairs of site indices and mutation types identifying the oncogenic variants in *g*. This approach enables estimation of mutation rates that are both gene-specific and context-dependent, accounting for trinucleotide mutational biases across the genome. The oncogenic mutation rates of the genes were consistently lower than the mutation rates of the genes.

Finally, the oncogenic mutation rates per somatic genotypes were computed by rescaling the per-gene background rates using the tumor mutation burden (TMB) observed in each genotype. Specifically, for a given cohort *c*, the mutation rate for a jump between the somatic genotype **x** the somatic genotype **y** is given by

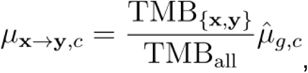

where TMB_{x,y},*c*_ is the average TMB in tumors with the somatic genotypes **x** or **y**, TMB_all_ is average TMB for all tumors, and *g* is the gene that is mutated in the somatic genotype **y** but not in **x** (the mutation gained by the jump). In cases where one genotype has no samples, the average is taken only from the observed partner. If both endpoints have zero samples, the rate defaults to the baseline per-gene rate 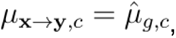 genotype classes. We considered TMB only from our largest dataset (TCGA) because after comparing the datasets with the Kolmogorov–Smirnov test, all of them proved to come from different distributions.

### Estimation of scaled selection coefficients and selective epistasis

To quantify the growth and survival advantages conferred by somatic SNVs in LUAD, we calculated scaled selection coefficients for recurrent mutations in a curated set of 1,200 genes [136,142–145]. These coefficients roughly equate to the ratio of the fixation rate for the oncogenic mutant (the rate at which the mutant is generated and subsequently selected to a high enough frequency to be observed as a somatic variant within the cancer cell population) to its background mutation rate. Precisely, we computed the fixation rate by analytical maximization of the binomial likelihood of observing at least one non-silent SNV in the gene given the number of samples with and without a non-silent SNV in the gene. Scaled selection coefficients for each mutated gene were estimated separately for ES- and NS-LUAD, as well as for a combined cohort. Within each group, we included samples with whole-exome, whole-genome, or targeted gene panel sequences (TGS). Results were consistent when excluding TGS samples, indicating robustness to sequencing strategy. Inclusion of TGS samples provided greater statistical power to detect significant differences in selection between cohorts. Results were also robust to cohort-specific normalization: applying median-scaling of coefficients prior to between-cohort comparison yielded consistent outcomes.

From the curated set of 1,200 genes, we selected 21 for detailed analysis of selective epistasis based on strong prior evidence as LUAD drivers [136,143] and their elevated average strength of selection across somatic genotypes. We quantified epistasis by calculating scaled selection coefficients conditional on the somatic genotype as in Alfaro-Murillo and Townsend [28]: for each pair or triplet of genes, we calculated the fixation rate of mutations in one gene conditional on the presence or absence of somatic variants in the others. We performed this calculation by maximizing the multinomial likelihood for the number of samples with each somatic genotype using PyMC [146]. The probability of a tumor being in a specific somatic genotype during the somatic evolutionary trajectory from organogenesis to biopsy was computed via a continuous-time Markov chain model [28]. We computed asymmetric 95% confidence intervals using Wilks’ theorem [147]. To obtain scaled selection coefficients for each mutation occurring within each somatic genotype, we divided the respective Poisson-corrected fixation rate [148] by the background oncogenic mutation rate of the gene.

To systematically assess selective epistasis in both the ever-smoker and the never-smoker groups, we evaluated 3,080 models representing all possible two- and three-gene combinations from the 21 LUAD driver genes. For each model, somatic genotypes were defined exclusively by the variant status (mutant or wildtype) of the genes included in each combination. To ensure complete genotype classification, we only included panel-sequenced samples in a given model if the sequencing panel covered all of the genes under analysis.

### Classification of synergistic and antagonistic selective epistasis

To classify a mutant somatic genotype as interacting synergistically or antagonistically with a certain mutation, we compared the scaled selection coefficients for a mutation when it occurred in the mutant somatic genotype to the scaled selection coefficients obtained for the same mutation in a somatic genotype lacking mutations in the genes included in the model. If the scaled selection coefficient for a mutation in gene *A* when it occurred in a wild-type genotype (∅ → *A*) was less than the scaled selection coefficient for a mutation in gene *A* when it occurred in a somatic genotype with a mutation in gene *B* (*B* → *A* + *B*), then we deemed the epistatic effect of mutant gene *B* on mutant gene *A* to be synergistic. If the scaled selection coefficient for ∅ → *A* was greater than the scaled selection coefficient for *B* → *A* + *B*, then we deemed the epistatic effect of mutant gene *B* on mutant gene *A* to be antagonistic. We further distinguished between two scales of antagonism: antagonistic sign epistasis when there was a reversal of the direction of selection—from positive (coefficient *γ* > 1) to negative selection (coefficient *γ* < 1)—in the antagonistic genotype, and antagonistic magnitude epistasis when there was a reduction in strength of selection but selection remained positive in the antagonistic genotype.

### Differential expression analysis

To evaluate the transcriptional impact of key driver mutations across smoking-stratified LUAD tumors, we performed differential expression analysis using RNA-seq data from the TCGA-LUAD project. We obtained unstranded raw read counts for 59 normal lung and 539 primary LUAD tumor samples using TCGAbiolinks v2.30.0. We excluded formalin-fixed paraffin-embedded samples and tumors which received neoadjuvant treatment to minimize technical and biological confounders. We retained solid-tissue normal samples as controls. We excluded genes with fewer than 10 total counts across all samples. For each sample, we assigned smoking status based on SBS4 signature weight from the matched MAF. The genes *EGFR*, *KRAS*, and *KEAP1* were considered mutated if at least one non-silent SNV, insertion, or deletion was present in the gene (excluding intronic regions). After this filtering and annotation, we validated that a principal-component analysis revealed no evidence of batch effects, confirming the suitability of the dataset for differential expression analysis.

We performed differential expression analysis with DESeq2 v1.42.1 [149] using two designs. Differential transcriptional effects of *EGFR* mutations in ES- and NS-LUAD were quantified as

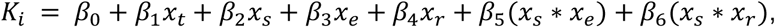

where *K*_*i*_ is the count for gene *i*, and *x*_*t*_, *x*_*s*_, *x*_*e*_, *x*_*r*_ are indicator variables for the sample type, smoking status, *EGFR* mutation status, and *KRAS* mutation status. The differential effect in question was measured by the interaction term between smoking status and *EGFR* allele status. The *EGFR*-focused-design accounts for the fact that most *EGFR*-wildtype LUADs are *KRAS*-mutant. Differential transcriptional effects of *KEAP1* mutations in ES- and NS-LUAD, were quantified as

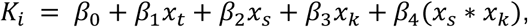

where *x*_*k*_ is the *KEAP1* mutation status.

We performed gene-set enrichment analysis with fgsea v1.28.0 using the MSigDB Hallmark gene sets. We ranked transcripts by signed *t*-statistics to capture both degree of significance and magnitude of fold change. We performed gene set variation analysis with GSVA v1.50.5 using the MSigDB canonical gene sets.

### Statistical Analysis

Differences in selection coefficients were assessed conservatively by non-overlapping 95% confidence intervals for each estimate. Differential gene expression was analyzed using two-tailed Wald tests, and statistical significance was determined based on Benjamini-Hochberg–adjusted *P* < 0.05. Gene set enrichment analyses were likewise considered significant at a Benjamini-Hochberg–adjusted *P* < 0.05. Analysis was performed in Python (3.9.5) and R (4.3.0).

## Data and code availability

All code, data, and instructions necessary to reproduce the results and figures in the current study is available in a public Zenodo repository, https://zenodo.org/records/16380037 [150] with no access restrictions.

## Acknowledgements

We thank Jeff Mandell for his advice on mutational signature attribution and mutation rate estimation, and Elizabeth Perry for assistance with gene expression analysis.

## Authors’ contributions

JPT, JAA-M, and KD conceptualized the project. KD curated the data and performed the formal analysis. KD and JAA-M wrote the software. KD, JAA-M, and JPT contributed to the methodology and investigation. KD drafted the original manuscript and JPT and JAA-M reviewed and edited the manuscript. All authors read and approved the final manuscripts.

## Materials and correspondence

Correspondence and material requests should be addressed to Jeffrey Townsend.

## Supporting information

**Supplementary Figure 1.**
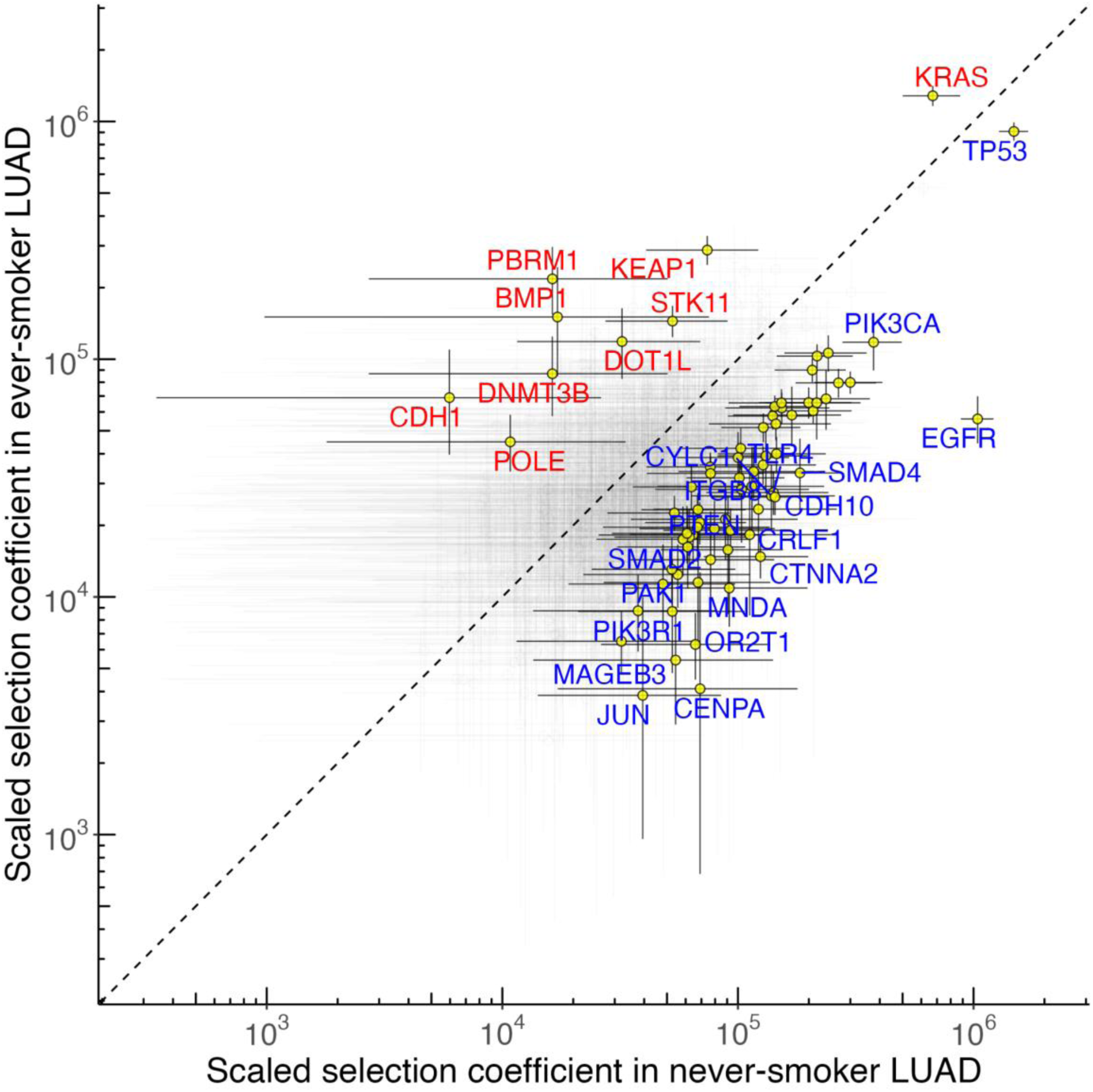
Strengths of somatic selection on 1,200 oncogenic gene mutations between lung adenocarcinoma (LUAD) in ever-smokers (*n* = 1,066) and in never-smokers (*n* = 656; significant: yellow circle, insignificant: no circle, red gene name: stronger selection in ES-LUAD, blue gene name: stronger selection in NS-LUAD).

**Supplementary Figure 2.**
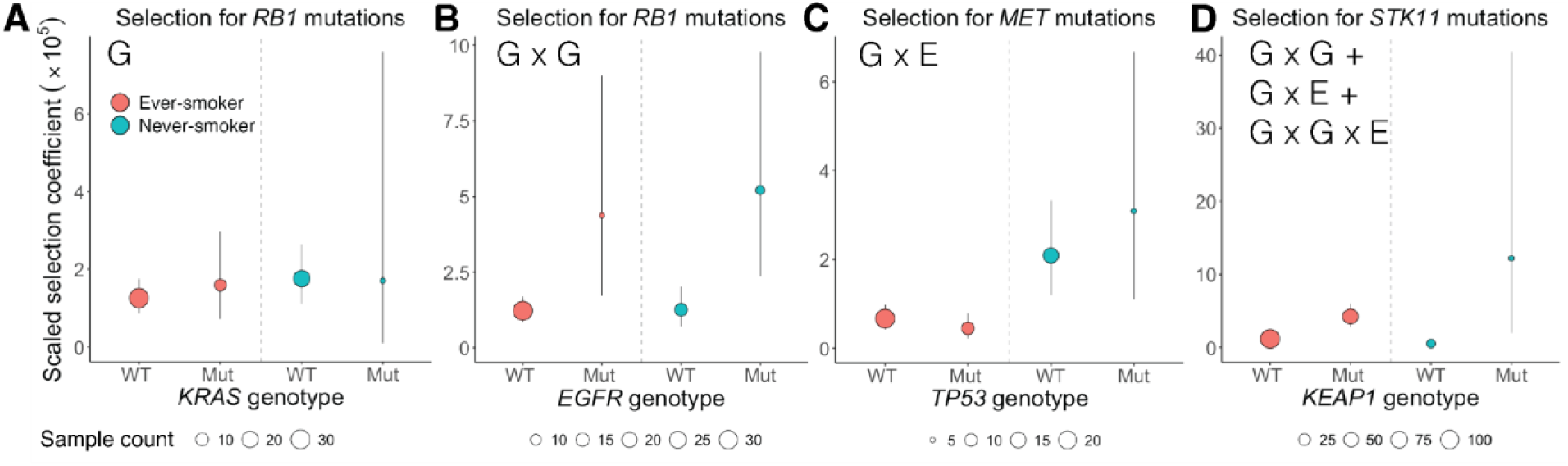
Four patterns of the influences of genetic and environmental interactions in ever-smoker and never-smoker lung adenocarcinoma. (**A**) *RB1* mutations with wild-type (WT) or mutant *KRAS*, (**B**) *RB1* mutations with wild-type or mutant *EGFR*, © MET mutations with wild-type or mutant *TP53*, and (**D**) *STk11* mutations with wild-type or mutant *KEAP1*.

**Table S1.**
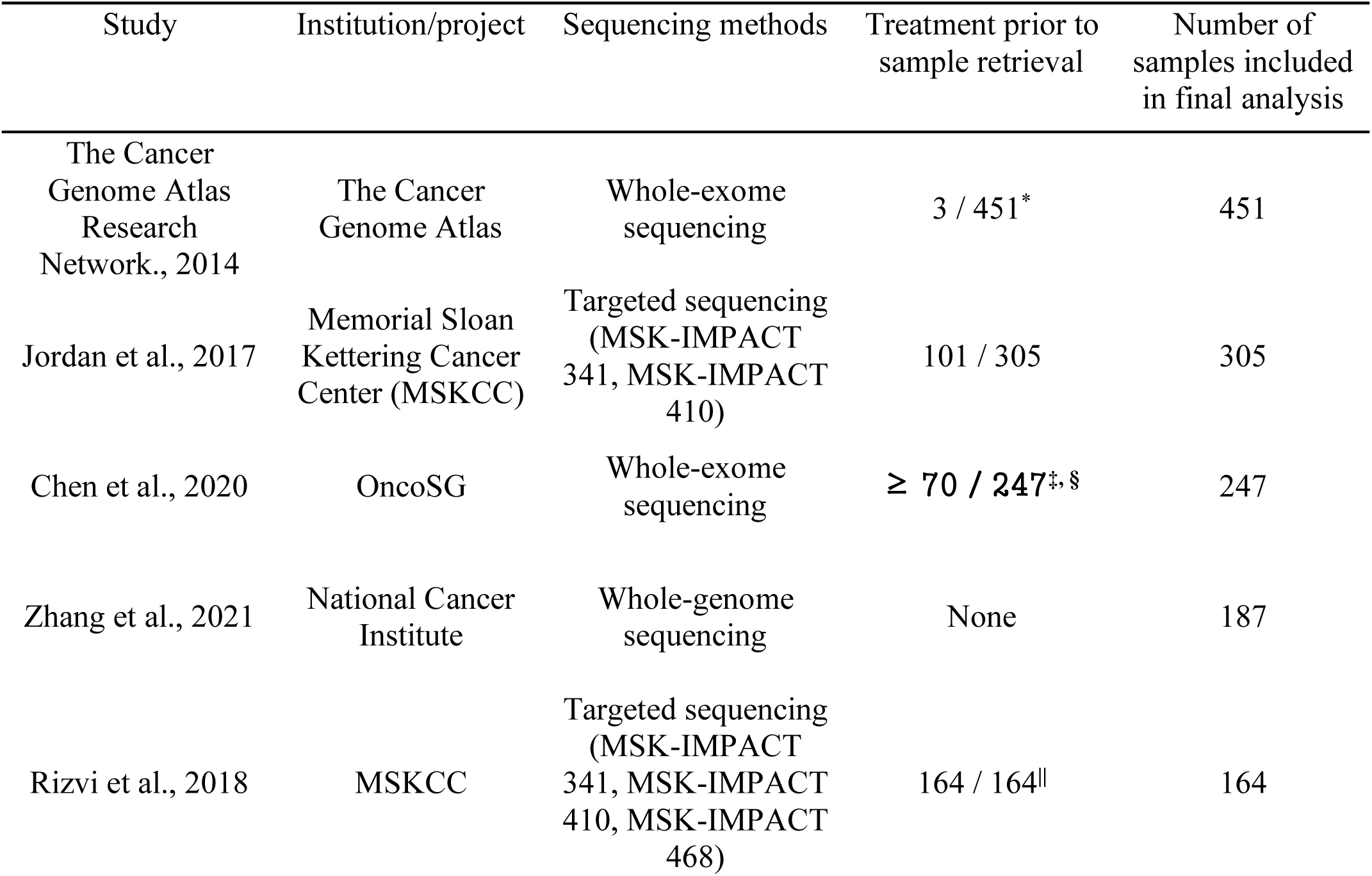
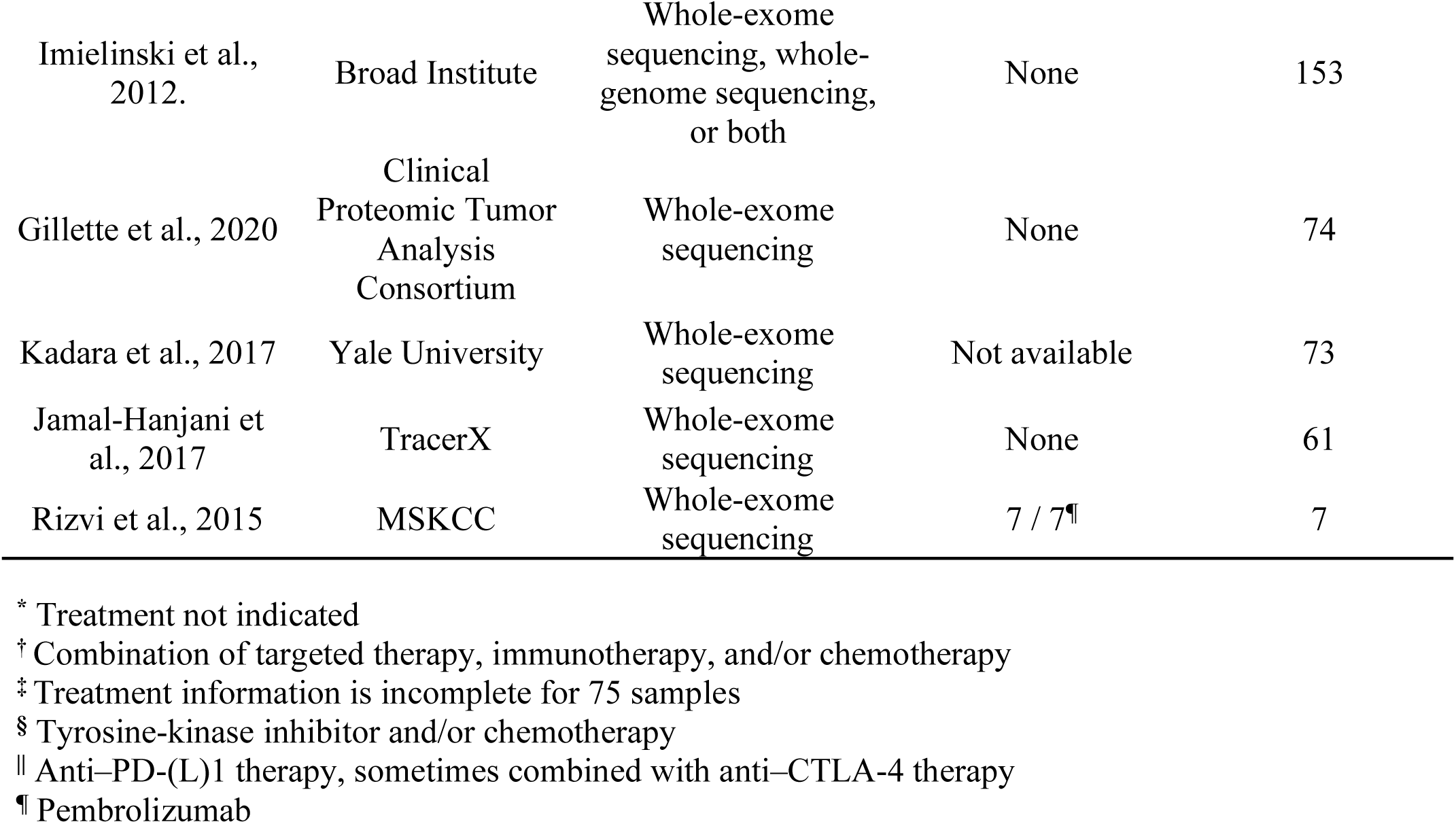
Characteristics of datasets included in study.

